## Supplementary Figures for "Ketamine evoked disruption of entorhinal and hippocampal spatial maps"

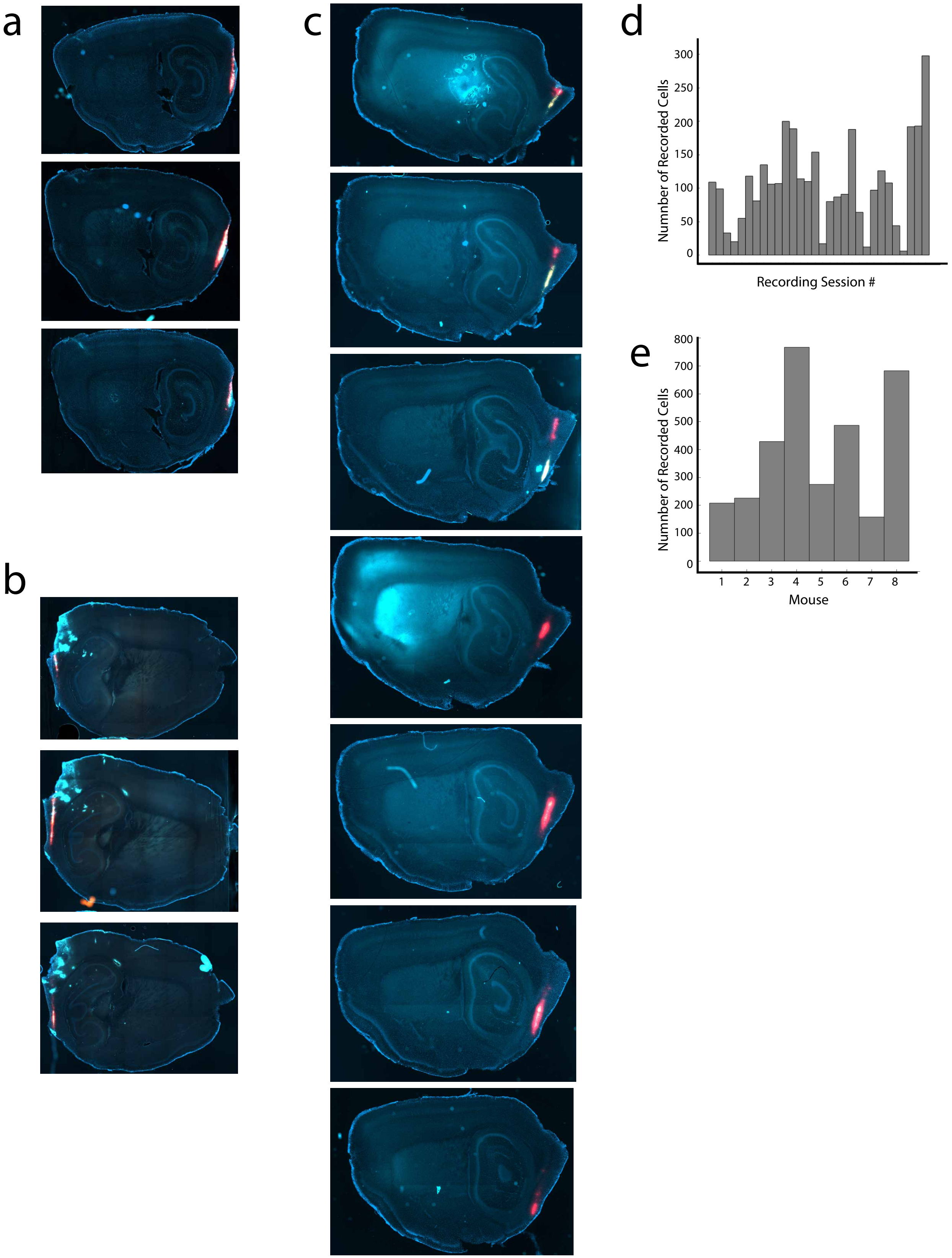
a
b
c
d
Numnber of Recorded Cells
300
250
200
150
100
50
0
Recording Session #
e
Numnber of Recorded Cells
800
700
600
500
400
300
200
100
0
1
2
3
4
5
6
7
8
Mouse

**Supplementary Figure 1**

a-c. Example histology of sagittal sections from three different mice (each sub panel corresponds to an individual mouse). Probe tracks are indicated by the fluorescent dye (red, yellow). Right hemisphere shown in (a), (c) and left hemisphere shown in (b).

d. Number of recorded cells labeled as good units by Kilosort 2.0 across all recording sessions.

e. Number of recorded cells labeled as good units by Kilosort 2.0 for each individual mouse (pooled across all recording sessions).

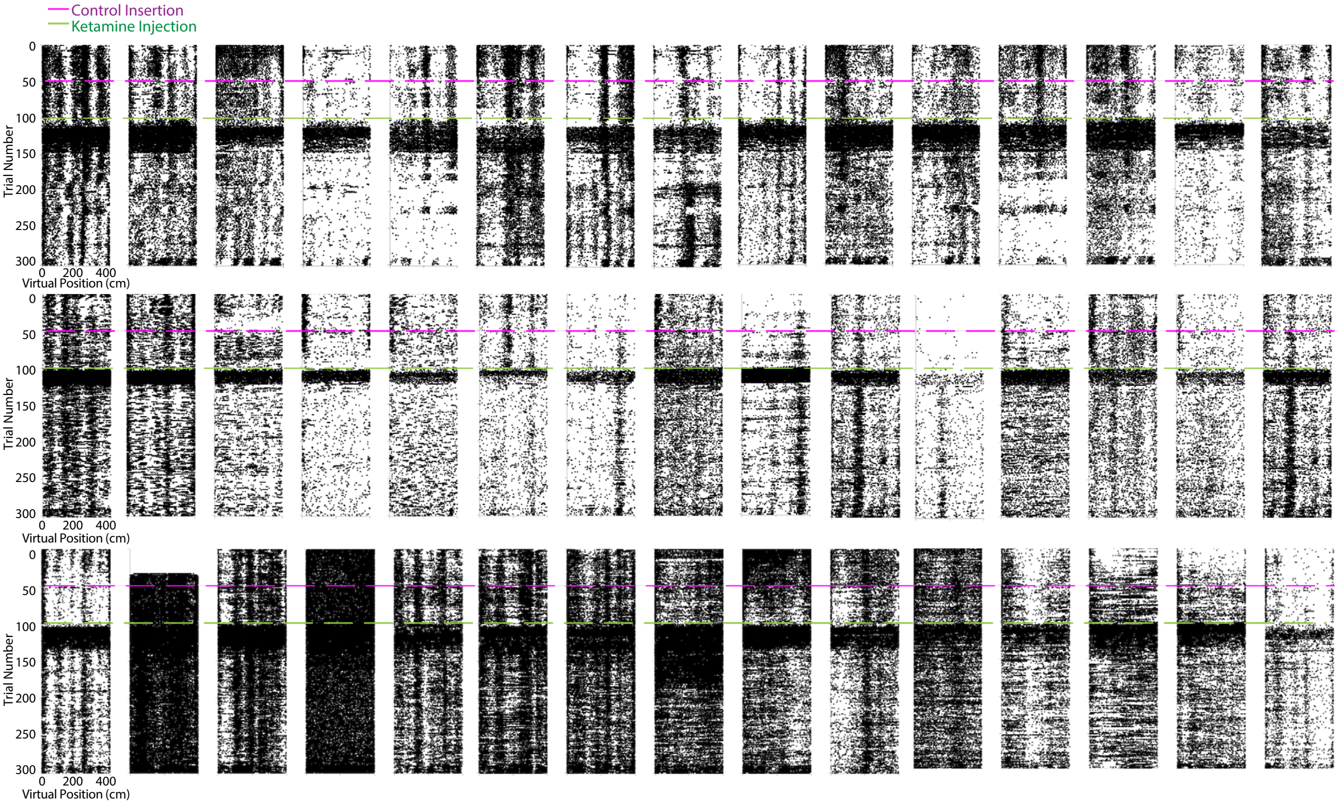

Control Insertion
Ketamine Injection
Trial Number
0
50
100
150
200
250
300
0 200 400
Virtual Position (cm)

**Supplementary Figure 2**

a. Spatial raster plots of 15 example cells from 3 different recording sessions in 3 different mice (each row is a unique session). Raster plots indicate individual spikes (black dots). The magenta line marks the beginning of the control epoch and the green line marks the beginning of the ketamine epoch.

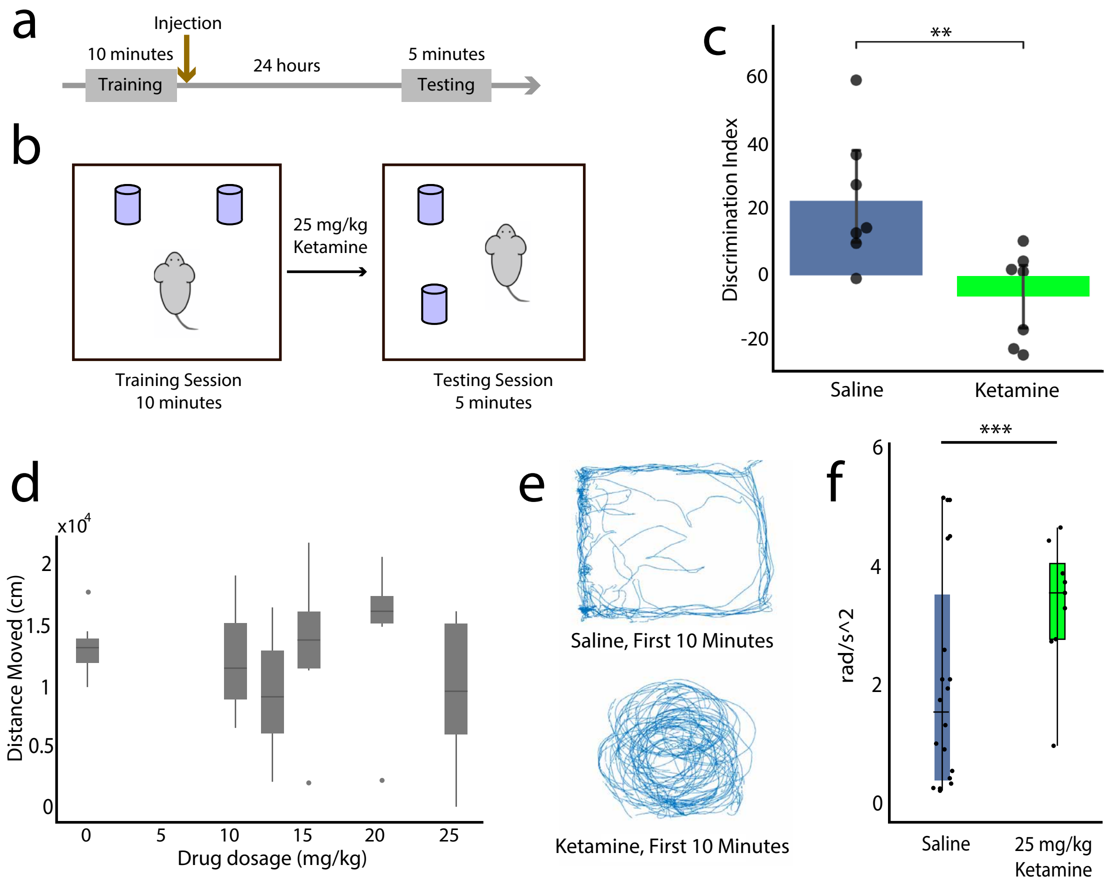
a
Injection
10 minutes
Training
24 hours
5 minutes
Testing
b
25 mg/kg
Ketamine
Training Session
10 minutes
Testing Session
5 minutes
c
**
Discrimination Index
60
40
20
0
-20
Saline
Ketamine
d
x10^4
Distance Moved (cm)
2
1.5
1
0.5
0
0
5
10
15
20
25
Drug dosage (mg/kg)
e
Saline, First 10 Minutes
Ketamine, First 10 Minutes
f
***
rad/s^2
6
4
2
0
Saline
25 mg/kg
Ketamine

**Supplementary Figure 3**

a. 14 mice performed an object-location memory task to test for spatial memory consolidation. In the training session, mice explored an open-field arena with two objects for 10 minutes. Immediately after the training session, half the mice were given 25 mg/kg ketamine and the other half were given a control saline injection. The following day, one of the objects was moved to a new spatial location and the mice were allowed to explore freely for 5 minutes.
b. Schematic of object-location memory tasks demonstrating how one object moved and one object did not move between the training and testing phase.
c. Discrimination index value for the 14 mice tested in the object-location memory task. The 7 mice that received the saline injection spent significantly more time investigating the object that moved, indicating preserved spatial memory consolidation (2-sided t-test $p=0.008$). Each black dot represents one mouse, the bar represents the mean and lines indicate SEM.
d. 5 male mice and 5 female mice were given ketamine at the following doses: 0 mg/kg, 10 mg/kg, 12.5 mg/kg, 15 mg/kg, 20 mg/kg, and 25 mg/kg. The drug was delivered to mice via an IP injection and the distance they moved in a 60 cm×60 cm open field was tracked for 30 minutes afterwards using Ethovision's behavior tracking software. Ketamine did not have consistent dose-dependent effects on median or interquartile range of distance moved.
e. Example trajectory of an individual mouse in a 60 cm×60 cm open field. cm open field arena over the 10 minutes following a saline control injection (top) and a 25 mg/kg ketamine injection (bottom). Mice that received a ketamine injection often displayed stereotyped spinning behaviors.
f. Mice that received ketamine had significantly increased angular acceleration in the 10 minutes when compared to the mice that received saline ($p<0.001$, 2-sided t-test) .

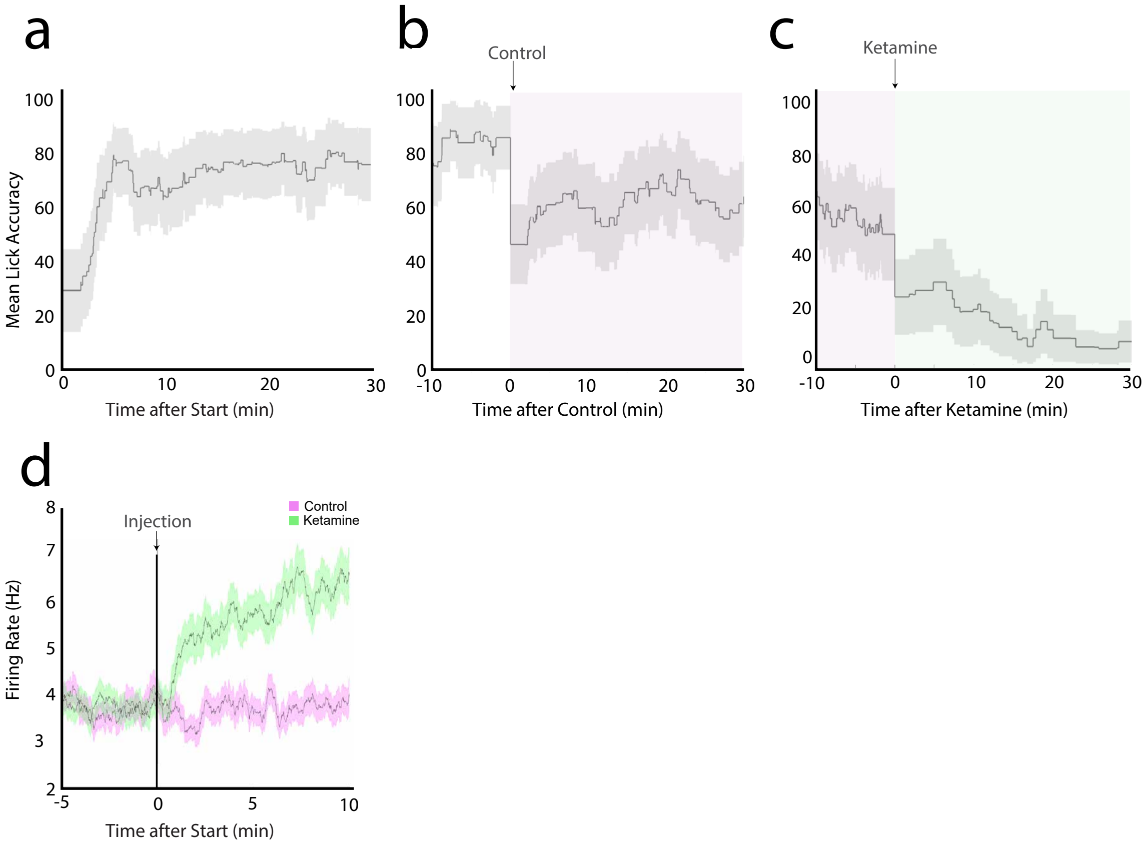
a
b
Control
c
Ketamine
Mean Lick Accuracy
100
80
60
40
20
0
0
10
20
30
Time after Start (min)
-10
Time after Control (min)
Time after Ketamine (min)
d
Injection
Control
Ketamine
Firing Rate (Hz)
8
7
6
5
4
3
2
-5
0
5
10
Time after Start (min)

**Supplementary Figure 4**

a. Mean mouse lick accuracy in the first 30 minutes of the baseline epoch (n = 32 sessions, 8 mice). Solid lines represent mean lick accuracy and shaded regions represent standard error of the mean.
b. Mean lick accuracy in the 10 minutes before the control injection and the 30 minutes after the control injection (n = 32 sessions, 8 mice). Solid lines and shading as reported in (a). Control epoch highlighted in magenta. Mean lick accuracy drops in the control epoch when examining accuracy over time but then remains stable.
c. Mean lick accuracy in the 10 minutes before the ketamine injection and 30 minutes after the ketamine injection (n = 32 sessions, 8 mice). Solid lines and shading as reported in (a). Control epoch highlighted in magenta and ketamine epoch highlighted in green. Mean lick accuracy drops significantly in the ketamine epoch when examining accuracy over time and approaches 0.
d. Mean firing rate of putative grid cells 5 minutes before and 10 minutes after either a control (magenta) or a 25 mg/kg ketamine (green) injection. Line indicates mean and shaded region shows SEM (n = 2135 cells, mouse n=8). Putative grid cells were identified by identifying cells whose firing rates were modulated by gain change.

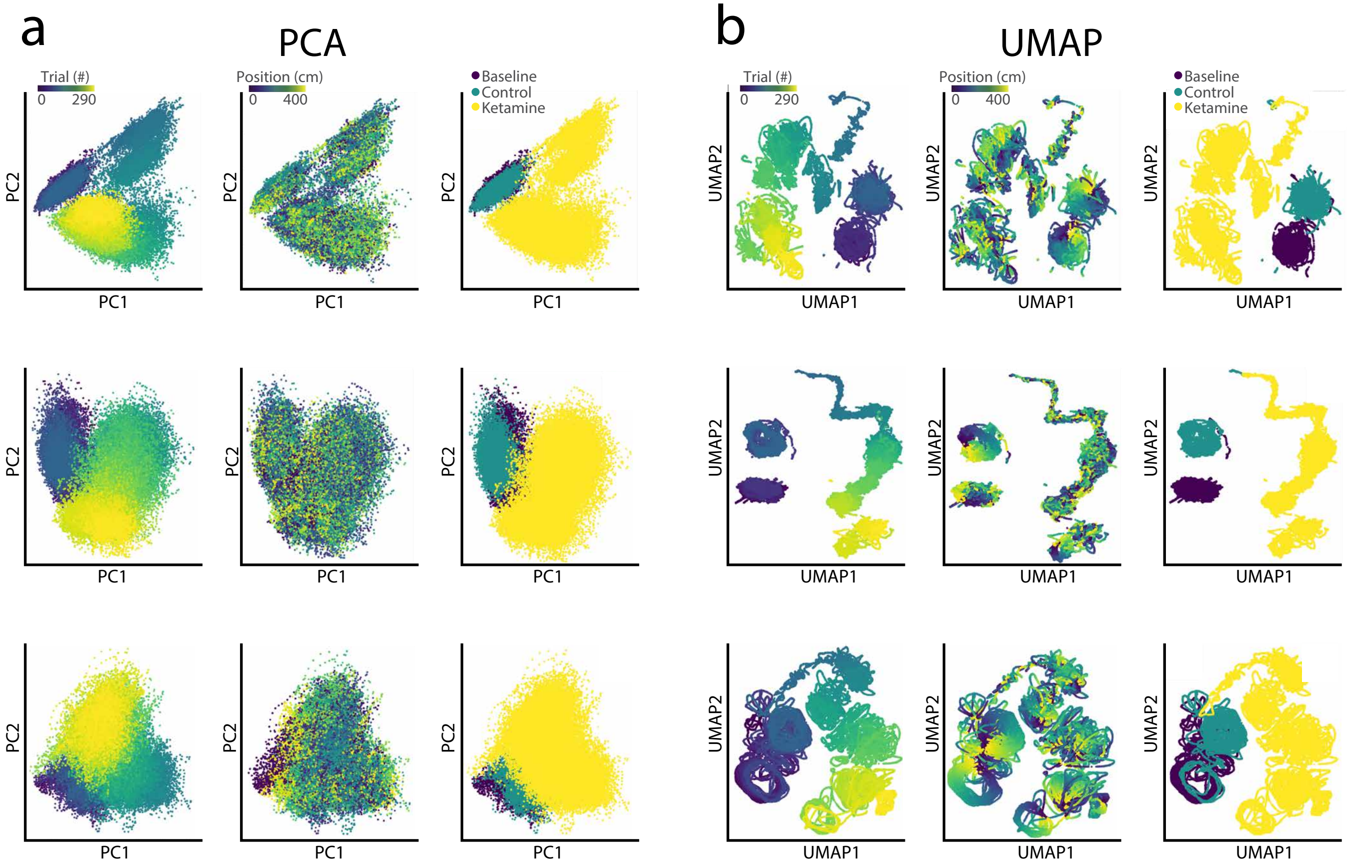
a
PCA
Trial (#)
0
290
Position (cm)
0
400
Baseline
Control
Ketamine
PC1
PC2
b
UMAP
UMAP1
UMAP2

**Supplementary Figure 5**

a. Three examples of temporally-binned firing rates of neural populations projected onto the first two PCA component dimensions. PCA dimensions colored by trial number (left column), VR position (middle column), and experimental epoch (right column).
b. Three examples of temporally-binned firing rates of neural populations projected onto the first two UMAP component dimensions. Colored by trial number (left column), VR position (middle column), and experimental epoch (right column).
